# D-GPM: a deep learning method for gene promoter methylation inference

**DOI:** 10.1101/438218

**Authors:** Xingxin Pan, Biao Liu, Xingzhao Wen, Yulu Liu, Xiuqing Zhang, Shengbin Li, Shuaicheng Li

## Abstract

**Background:** Gene promoter methylation plays a critical role in a wide range of biological processes, such as transcriptional expression, gene imprinting, X chromosome inactivation, *etc*. Whole-genome bisulfite sequencing generates a comprehensive profiling of the gene methylation levels but is limited by a high cost. Recent studies have partitioned the genes into landmark genes and target genes and suggested that the landmark gene expression levels capture adequate information to reconstruct the target gene expression levels. Moreover, the methylation level of the promoter is usually negatively correlated with its corresponding gene expression. This result inspired us to propose that the methylation level of the promoters might be adequate to reconstruct the promoter methylation level of target genes, which would eventually reduce the cost of promoter methylation profiling.

**Results:** Here, we developed a deep learning model (D-GPM) to predict the whole-genome promoter methylation level based on the methylation profile of the landmark genes. We benchmarked D-GPM against three machine learning methods, namely, linear regression (LR), regression tree (RT) and support vector machine (SVM), based on two criteria: the mean absolute deviation (MAE) and the Pearson correlation coefficient (PCC). After profiling the methylation beta value (MBV) dataset from the TCGA, with respect to MAE and PCC, we found that D-GPM outperforms LR by 9.59% and 4.34%, RT by 27.58% and 22.96% and SVM by 6.14% and 3.07% on average, respectively. For the number of better-predicted genes, D-GPM outperforms LR in 92.65% and 91.00%, RT in 95.66% and 98.25% and SVM in 85.49% and 81.56% of the target genes.

**Conclusions:** D-GPM acquires the least overall MAE and the highest overall PCC on MBV-te compared to LR, RT, and SVM. For a genewise comparative analysis, D-GPM outperforms LR, RT, and SVM in an overwhelming majority of the target genes, with respect to the MAE and PCC. Most importantly, D-GPM predominates among the other models in predicting a majority of the target genes according to the model distribution of the least MAE and the highest PCC for the target genes.

## 1 Background

By influencing the transcription factor’s accessibility to DNA, the methylation of the promoter of a gene regulates various biological process [1]. Enzymatic digestion, affinity enrichment, and bisulfite conversion are methods to capture the DNA methylation level [2]. Despite technological advances, there are still limitations in existing wet-lab methods. The resolution of enzymatic digestion-based approach is restricted to regions adjacent to the methylation-sensitive restriction enzyme recognition sites [3]. The resolution of methylated DNA immunoprecipitation is limited to 100-300 base pair long fragments, and it is also biased towards the hypermethylated regions[4]. Illumina’s 450K bead-chip is the most widely used method for profiling DNA methylation in humans, but the chip only probes approximately 450K CpG sites in the human genome and covers partial CpG islands and may be biased towards the CpG dense contexts [5]. Whole-genome bisulfite sequencing is a golden standard protocol. However, it is too costly because the genome-wide deep sequencing of bisulfite-treated fragments needs to generate a compendium of gene methylation level over a large number of conditions, such as retrovirus, DNMT activity changes and drug treatments [6]. The community awaits more feasible and more economical solutions.

Previous research suggests that there is a low-rank structure in the genome-wide gene expression profile [7], i.e., by leveraging the inner correlation between genes, the expression level of a few well-chosen landmark genes captures sufficient detail to reconstruct the expression of the rest of genes, namely, the target genes across the genome. The above result was achieved by studying the gene regulation networks and conducting a principal component analysis on the whole-genome expression profile [8]. Consequently, scientists created a new technology called L1000, which only acquires the expression profiling of the landmark genes (~1000) to infer the expression profiling of the target genes (~21000) [7].

Inspired by L1000, we proposed a method according to the following rationale to acquire the whole-genome promoter methylation level according to the promoter mythylations of the landmark genes. First, latent associations exist between the expression of these landmark genes and the target genes at the genome-wide level [9]; second, the methylation in the promoters located upstream of the transcription start site is usually negatively correlated with their corresponding gene expression levels [10]. Hence, it is likely that strong associations are present among the methylation levels in the landmark genes and target genes.

To predict the methylation panorama on the whole genome is a large-scale multitask machine learning problem, with a high-dimensional aim (~21,000) and a low-dimensional attribute (~1,000). Meanwhile, the deep learning method has shown its power in integrating large-scale data and capturing the nonlinear complexity of input features. In biology, extensive applications include predictions for the splicing activity of individual exons, inferring chromatin marks from the DNA sequence, and the quantification of the effect of SNVs on chromatin accessibility [11-13].

Here, we present a multilayer deep neural network named Deep-Gene Promoter Methylation Inference (D-GPM). To evaluate our D-GPM model, we benchmarked its performance against linear regression (LR), regression tree (RT) and support vector machine (SVM), with regard to methylation profile data based on the Illumina Human Methylation 450k data from The Cancer Genome Atlas (TCGA) [14]. LR is used to infer the methylation levels of the target genes based on the promoter methylation of the landmark genes using linear regression models. However, the linear model may fail to capture the nonlinear relations of the original data. The SVM reliably represents complex nonlinear patterns [15] but is limited by poor scalability due to the large data size. The RT addresses the interpretability of the biological data and prediction model despite its lower accuracy and instability in some predictors.

According to Illumina Human Methylation 450k data, we accessed the methylation information from 902 landmark genes and 21645 target genes. The experimental results show that D-GPM consistently outperforms the other methods for the testing data, and this was measured using the criteria of the mean absolute deviation (MAE) and the Pearson correlation coefficient (PCC).

## 2 Results

Here, we introduced the methylation beta value (MBV) datasets from TCGA and defined the methylation profile inferences as a multitask regression problem, with the MBV-tr for training, MBV-va for validation and MBV-te for testing. We have also illustrated our deep learning method D-PGM and the other three methods, including LR, RT and SVM, to work out the regression problem. Next, we show the predictive performances of the above methods on the MBV-te data based on the MAE and PCC criteria.

### 2.1 D-GPM performs the best for predicting promoter methylation

The back-propagation algorithm, mini-batch gradient descent, and other beneficial deep learning techniques are adopted in training the D-GPM [16]. The detailed parameter configurations are shown in Table 1.

**Table 1.** Detailed parameter configurations are given in D-GPM.

| Parameters |  |
| --- | --- |
| # of hidden layers | [1-8] |
| # of hidden units in each hidden layer | [1000,2000,3000,4000,5000,6000,7000,8000,9000] |
| Dropout rate | [0%,5%,10%,15%,20%,25%,30%,35%,40%,45%,50%] |
| Momentum coefficient | 0.5 |
| Initial learning rate | 5E-4, 2E-4, 1E-4 or 8E-5 |
| Minimum learning rate | 1.00E-05 |
| Learning rate decay factor | 0.9 |
| Learning scale | 3.0 |
| Mini-batch size | 200 |
| Training epoch | 500 |
| Weights initial range | $\left[ -\frac{\sqrt{6}}{\sqrt{n_i + n_o}}, \frac{\sqrt{6}}{\sqrt{n_i + n_o}} \right]$ |

According to the parameter configurations, all the combinations of parameters are made during the D-GPM training for predicting the promoter methylation of the target genes.

As Table 2 indicates, D-GPM acquires the best MAE performance on MBV-te, with five hidden layers of 7000 units and a 15% dropout rate (D-GPM-15%-7000×5) among the 792 (8*9*11) various D-GPMs. Meanwhile, D-GPM has an extraordinary edge over MAE compared with LR, SVM, and RT.

**Table 2.**
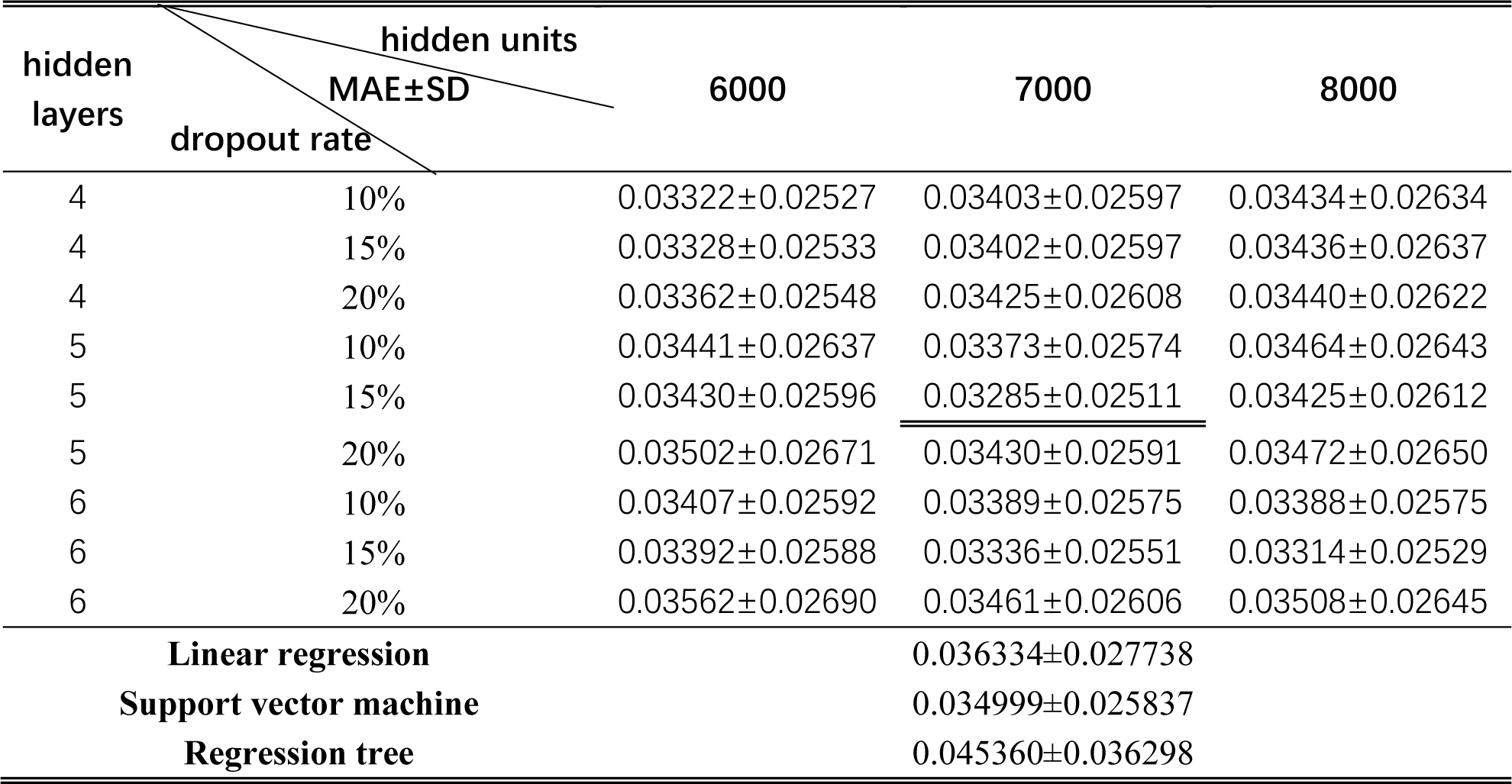
The MAE-based overall errors of LR, RT, SVM, and D-GPM, with partially different architectures and partially different dropout rates on MBV-te. The numbers after “±” are the standard deviations of the prediction errors over all the target genes. The best performance of D-GPM is underlined. SD refers to standard deviation.

Similarly, D-GPM also obtains the best PCC performance on MBV-te among the 792 prediction models, as shown in Table 3. The complete MAE, PCC evaluation of D-GPM armed with the other architectures (hidden layer: from 1 to 8, with step size 1; hidden unit: from 1000 to 9000, with step size 1000; dropout rate: from 0% to 50%, with step size 5%) on the MBV-te is given in the Supplementary Material.

**Table 3.**
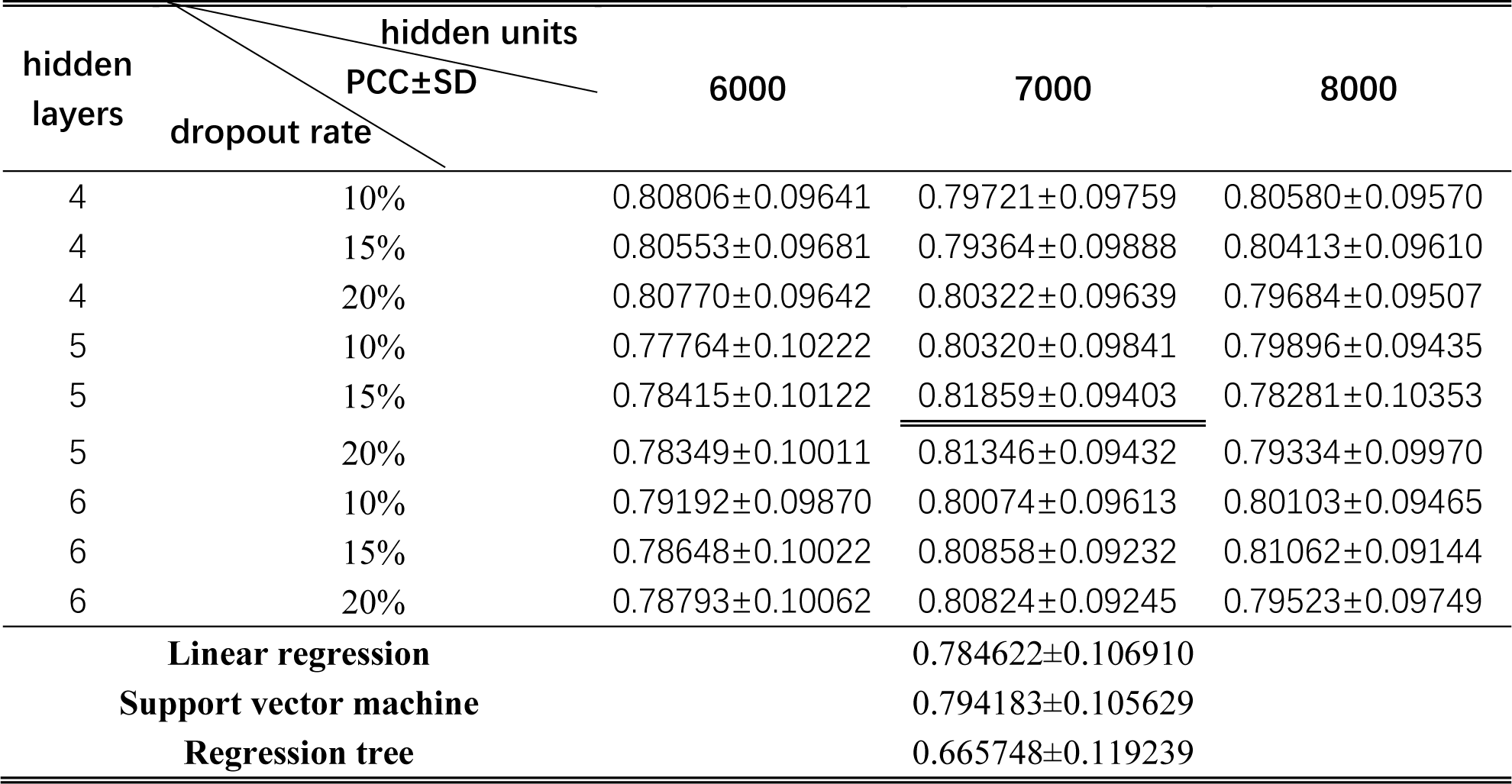
The PCC of LR, RT, SVM and D-GPM, with partially different architectures and partially different dropout rates on MBV-te. The numbers after “±” are the standard deviations of the PCC over all the target genes. The best performance of D-GPM is underlined. SD refers to standard deviation.

Based on the above MAE and PCC, we can conclude that D-GPM is the best model for predicting promoter methylation among the prediction models.

### 2.2 Evaluation according to MAE criteria

D-GPM acquires the best MAE performance on the MBV-te, with a 15% dropout rate (described as D-GPM-15%) among the eleven dropout rates, ranging from 0% to 50%, with a step size of 5%. Fig. 1 shows the overall MAE performances of D-GPM-15% and SVM on the MBV-te. The larger architecture of D-GPM-15% (five hidden layers with 7000 hidden units in each hidden layer, described as D-GPM-15%-7000×5) acquires the least MAE on the MBV-te. The improvements of D-GPM-15%-7000×5 is 9.59%, 27.58%, and 6.14% over LR, RT, and SVM, respectively. A possible explanation is that deep learning can capture complex features [17].

**Fig. 1.**
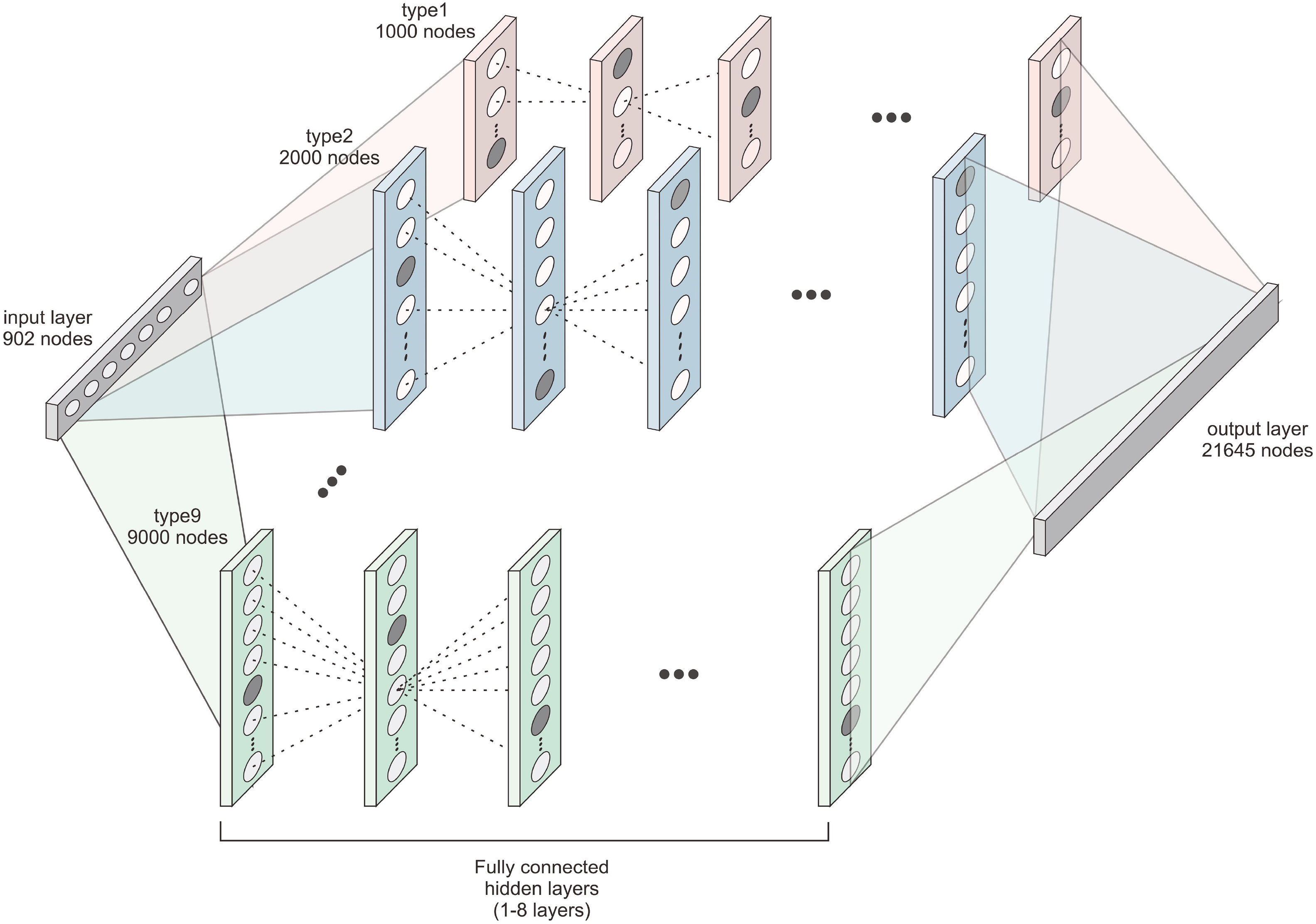
The overall MAE errors of D-GPM-15%, with various architectures on the MBV-te.

D-GPM also outperforms LR and RT for almost all the target genes on MAE. Fig. 2 shows the density plots of the MAE errors of all the target genes by LR, RT, SVM, and D-GPM. On the whole, D-GPM occupies a larger proportion at the low MAE level and a lower proportion at the high MAE level compared to the three machine learning methods, especially for RT and LR, attesting to the prominent performance of D-GPM.

**Fig. 2.**
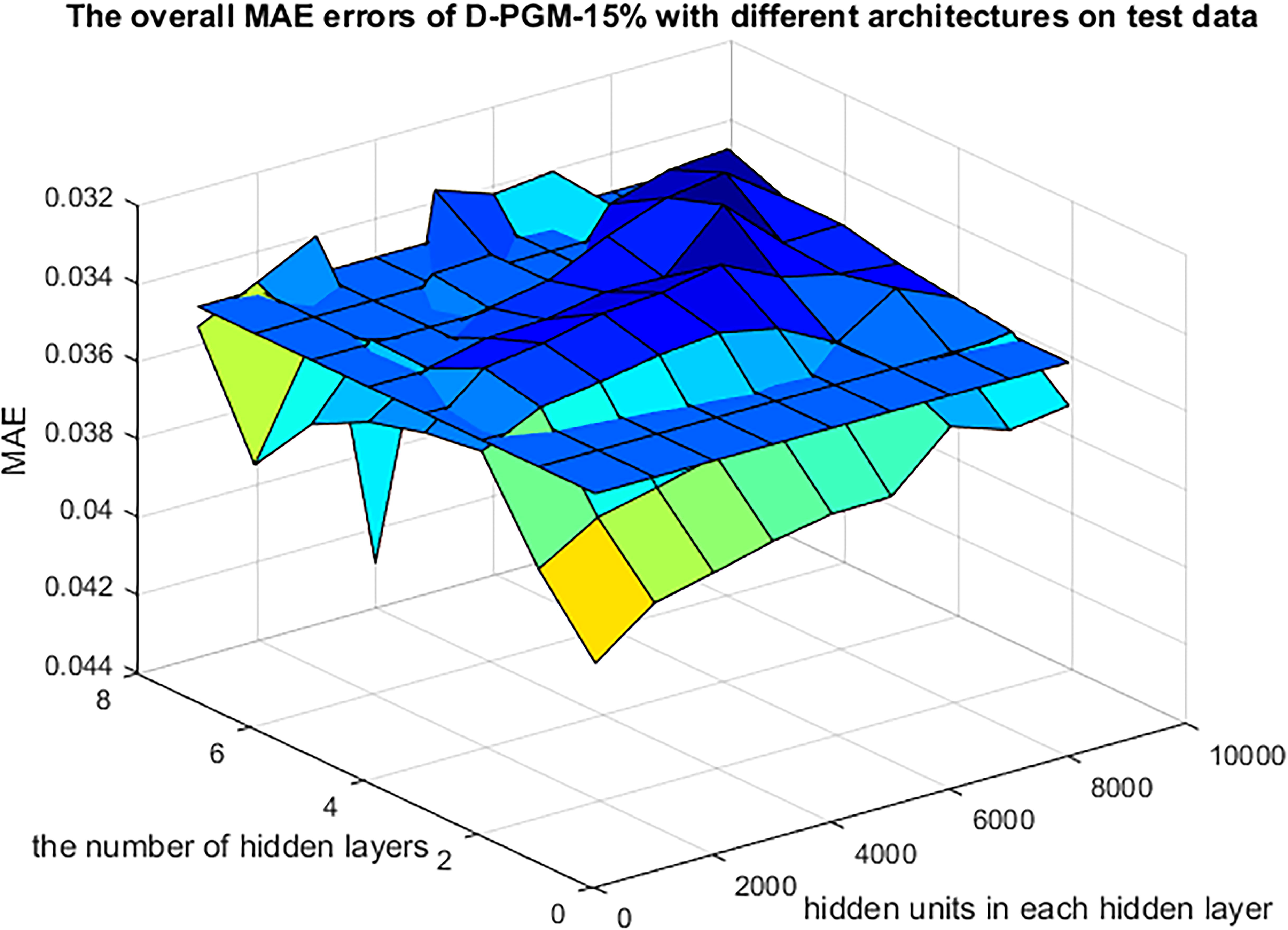
The density plots of the MAE by LR, RT, SVM, and D-GPM on the MBV-te.

Fig. 3(a-c) displays a genewise comparative analysis between D-GPM and the other three methods. In terms of MAE, D-GPM outperforms RT for 95.66% (20705 genes) of the target genes, it outperforms LR for 92.65% (20054 genes) of the target genes, and it outperforms SVM for 85.49% (18504 genes) of the target genes. These results can also be viewed by the larger proportion of dots lie above the diagonal, and this better performance may suggest that D-GPM can capture some intrinsic nonlinear features of the MBV data, which LR, RT, and SVM did not accomplish.

**Fig. 3.**
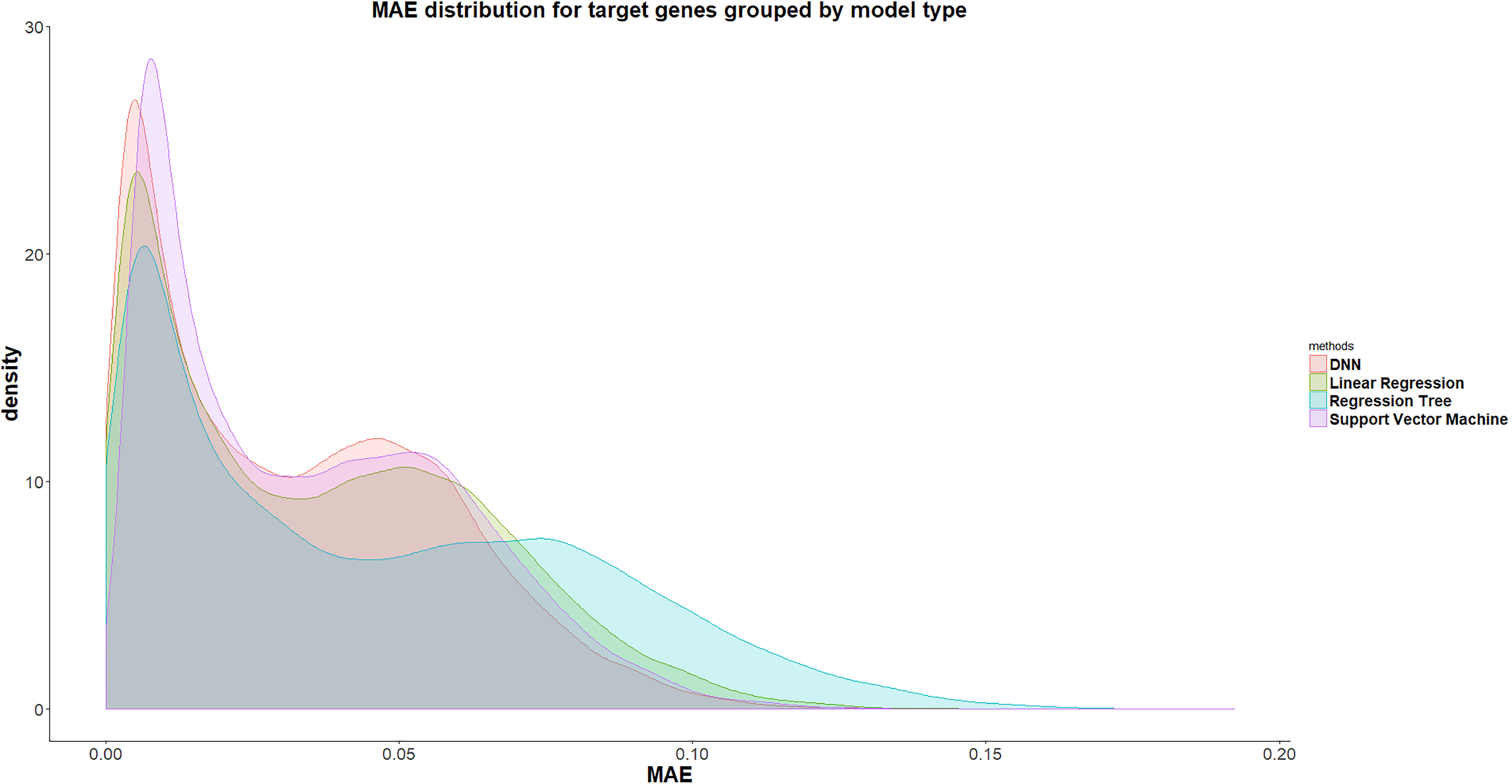
(a) The MAE errors of each target gene by D-GPM compared with RT on the MBV-te. (b) The MAE errors of each target gene by D-GPM compared with LR on the MBV-te. (c) The MAE errors of each target gene by D-GPM compared with SVM on the MBV-te. Among (a), (b) and (c), each dot represents one out of the 21645 target genes. The x-axis is the MAE of each target gene by D-GPM, and the y-axis is the MAE of each target gene by the other machine learning method. The dots above the diagonal indicate that D-GPM achieves lower error compared with the other method.

RT performs significantly worse than the other methods in the MAE aspect. One possible reason is that the model is oversimplified to capture the essential features between the landmark and target genes based on the MBV-te [18].

According to the model distribution of the lowest MAE for each target gene, we find the best model distribution, as shown in Fig. 4(a). RT accomplishes the best MAE performance for 305 target genes (1.41%), including the genes *BRD2, GPI, MAF*, and *MICB*, implying that there is a relatively simple methylation regulation mechanism and promoter methylation of these genes that is dominantly regulated by a very few landmark genes. The LR can predict 1,242 target genes (5.74%), at best, among the other three methods, including the genes *ABCD1, HPD, AMH* and *ARAF*, laying a solid foundation for the pathogenesis of diseases, such as Adrenoleukodystrophy, Hawkinsinuria, Persistent Mullerian Duct Syndrome and Pallister-Killian Syndrome, using our LR [19-22]. Noticeably, SVM performs best for a total of 2813 genes (13.00%), including *ACE2, A2M*, and *CA1*. One possible explanation for this is that there seem to be intricate and complicated interactions among the promoter methylation of the landmark genes and these 2813 target genes. Undoubtedly, D-GPM does better than the other three methods, as far as 17,285 genes (79.86%) are concerned, demonstrating the deep neural networks’ powerful ability to capture the nonlinear relationship of methylation profiling.

**Fig. 4.**
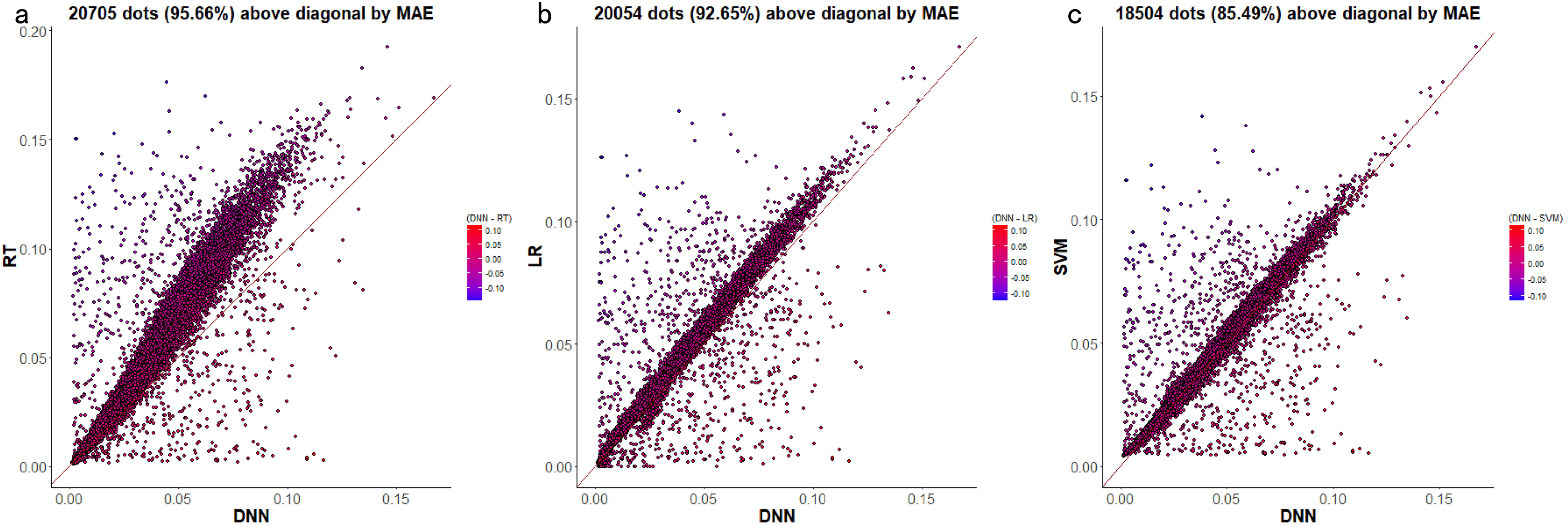
(a) Distribution of the best model according to MAE for the target genes. (b) Distribution of best model according to PCC for the target genes.

### 2.3 Evaluation according to PCC

D-GPM accomplishes the best PCC performance on the MBV-te, with a 15% dropout rate. Fig. 5 shows the overall PCC performances of D-GPM-15% and the other methods on MBV-te. Similar to the MAE, D-GPM-15%-7000×5 acquires the most significant PCC on the MBV-te. The relative improvement of D-GPM-15%-7000×5 is 4.34% over the LR, 22.96% over RT and 3.07% over SVM. Similar to MAE, almost all the combined architectures of D-GPM-15% outperforms LR and RT in PCC performance.

**Fig. 5.**
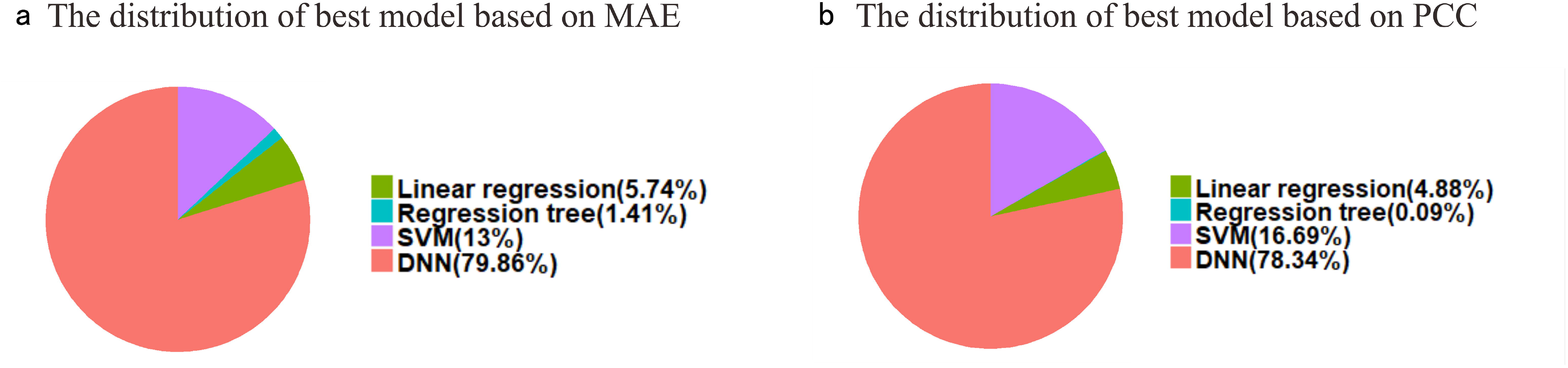
The overall PCC performance of D-GPM-15% with various architectures on the MBV-te.

In terms of the PCC, D-GPM also outperforms RT and LR for almost all the target genes. Fig. 6 displays the density plots of the PCC of all the target genes by LR, RT, SVM, and D-GPM. By and large, we can see that D-GPM possesses a larger proportion of the high PCC and a lower proportion at the low PCC compared to the RT, LR, and SVM.

**Fig. 6.**
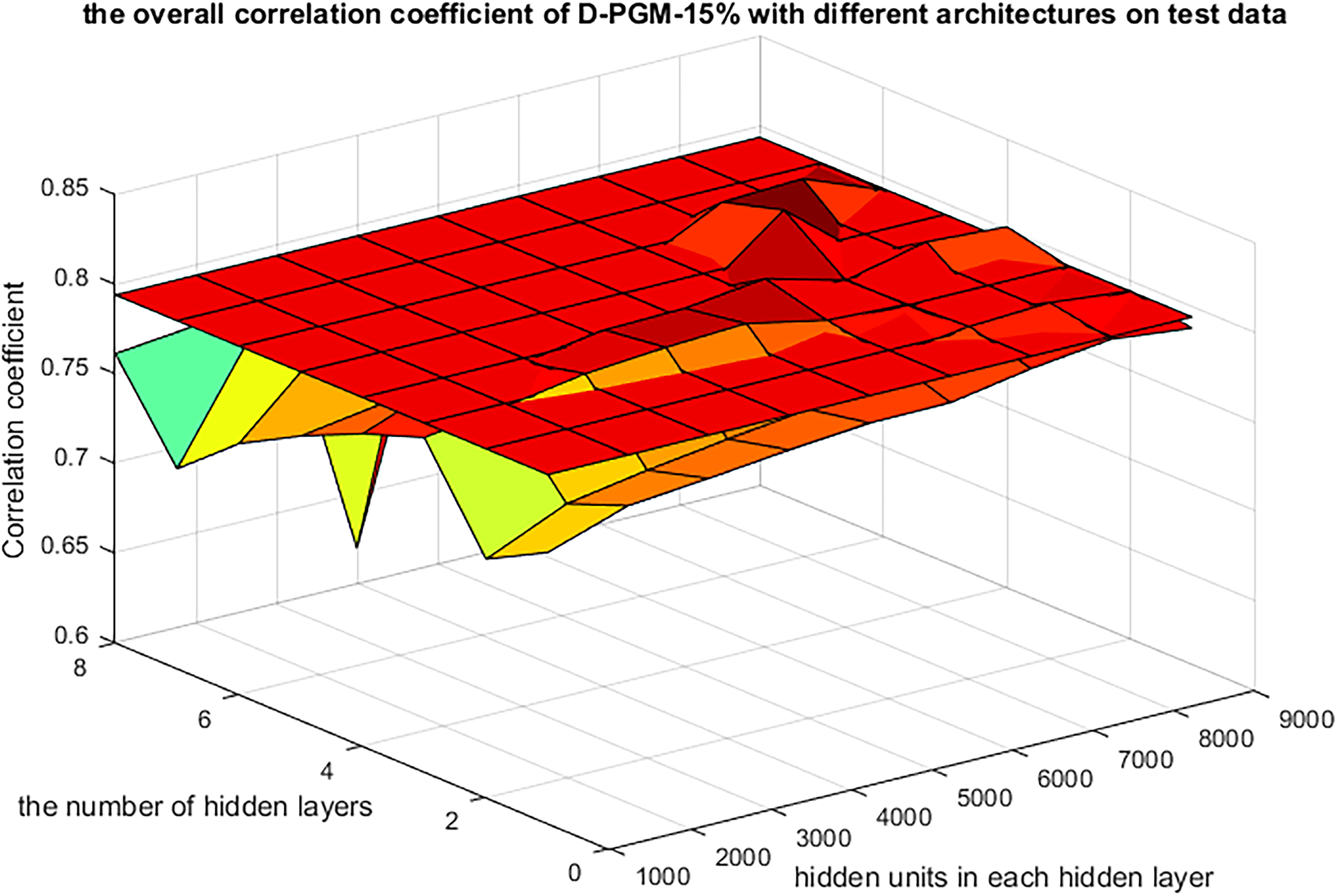
The density plots of the PCC of all the target genes by LR, RT, SVM and D-GPM on MBV-te.

Fig. 7(a-c) shows a genewise comparative analysis between D-GPM and the other three methods. For PCC, D-GPM outperforms RT in 98.25% (21266 genes) of the target genes, outperforms LR in 91.00% (19696 genes) in of the target genes and outperforms SVM in 81.56% (17653 genes) of the target genes. Therefore, D-GPM’s powerful prediction for the PCC of the overall target genes is preserved similar to its effective prediction for the MAE. It is obvious that although the prediction property of the SVM is still modest in the PCC aspect, its PCC for some of the target genes is significantly higher than D-GPM and behaves better than the prediction for the same target genes in the MAE aspect. This finding is probably due to the fact that SVM is based on the principle of the structural risk minimization, avoiding overlearning problems and having a strong generalization ability.

**Fig. 7.**
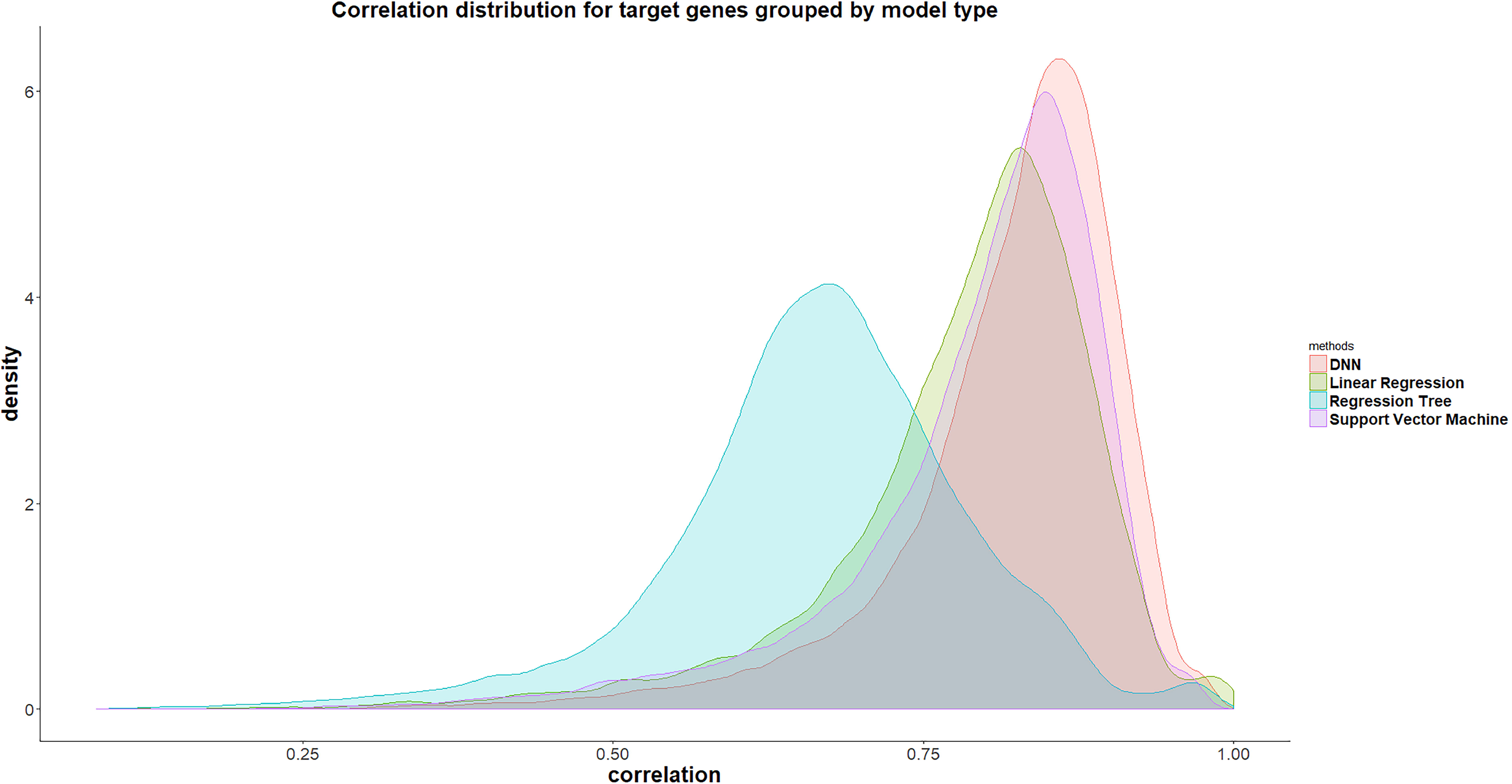
(a) The PCC performance of each target gene by D-GPM compared with RT on the MBV-te. (b) The PCC performance of each target gene by D-GPM compared with LR on the MBV-te. (c) The PCC performance of each target gene by D-GPM compared with SVM on the MBV-te. Among (a), (b) and (c), each dot represents one out of the 21645 target genes. The x-axis is the PCC of each target gene by the abovementioned three machine learning technique, and the y-axis is the PCC of each target gene by D-GPM. The dots above the diagonal indicate that D-GPM achieves high PCC compared with the other method.

According to the model distribution of the maximal PCC, we find the best model distribution, as shown in Fig. 5(b). Surprisingly, RT only gains the best PCC performance for 19 target genes (0.09%), including the genes *ALG1* and *NBR2* falling far below the best 305 target genes in the MAE aspect. Considering RT’s awful prediction power for PCC compared with its better prediction power for MAE, it may be explained by the fact that RT makes decisions based on an over simple assumption. LR predicts the best 1057 target genes (4.88%) among other three methods, including the genes *AASS* and *ACE*, which is almost the same as the 1242 in the MAE level. For PCC, SVM is on its best behavior for a total of 3613 genes (16.69%), having an increasing number compared to that of the MAE, in contrast to RT. Undoubtedly, D-GPM outperforms the other three methods, with regard to 16956 genes (78.34%), but behaves badly relative to the 79.86% in the MAE level, suggesting an inability to predict the PCC for the target genes.

Noticeably, the dropout regularization manages to improve the performance of D-GPM-7000×5 on the MBV-te, as shown in Fig. 8. With a 15% dropout rate, D-GPM-15%-7000×5 consistently achieves the best MAE performance on the MBV-te among the models, with 0%, 15%, 20%, 25%, and 35% dropout rates, proving that overfitting and underfitting both result in a bad influence on the prediction model.

**Fig. 8.**
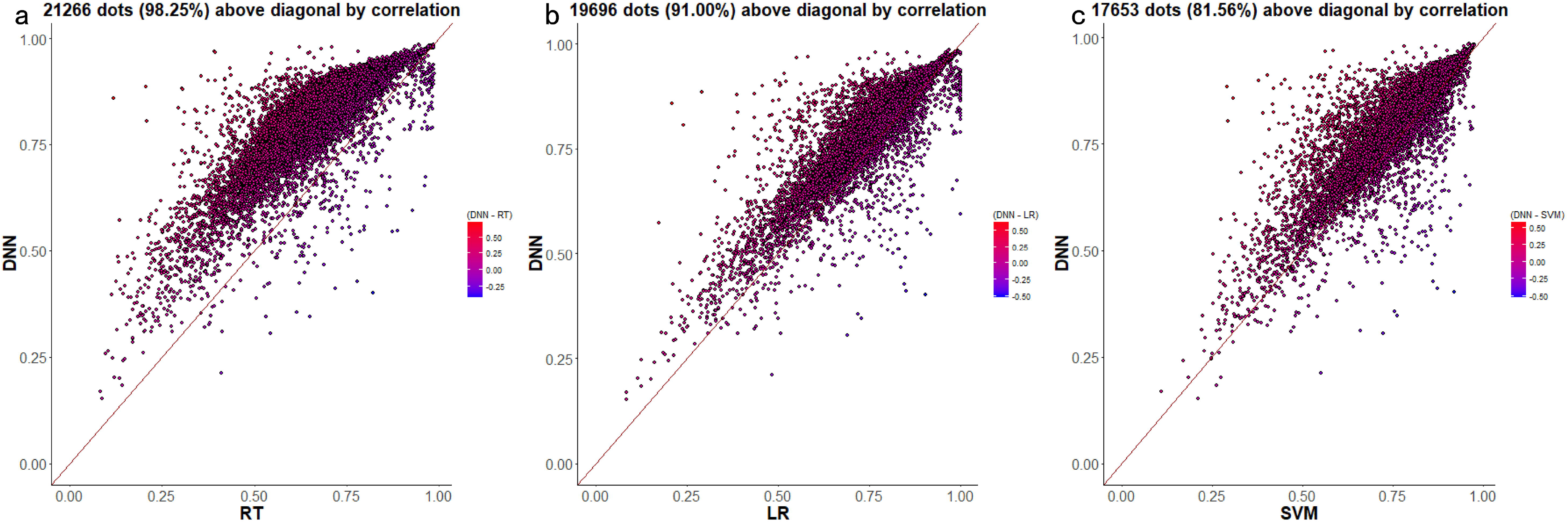
The overall MAE decreasing the curves of D-GPM-7000×5 on the MBV-te, with different dropout rates. The x-axis is the training epoch, and the y-axis is the overall MAE error. The overall MAE error of SVM is also covered for comparison.

**Fig. 9.**
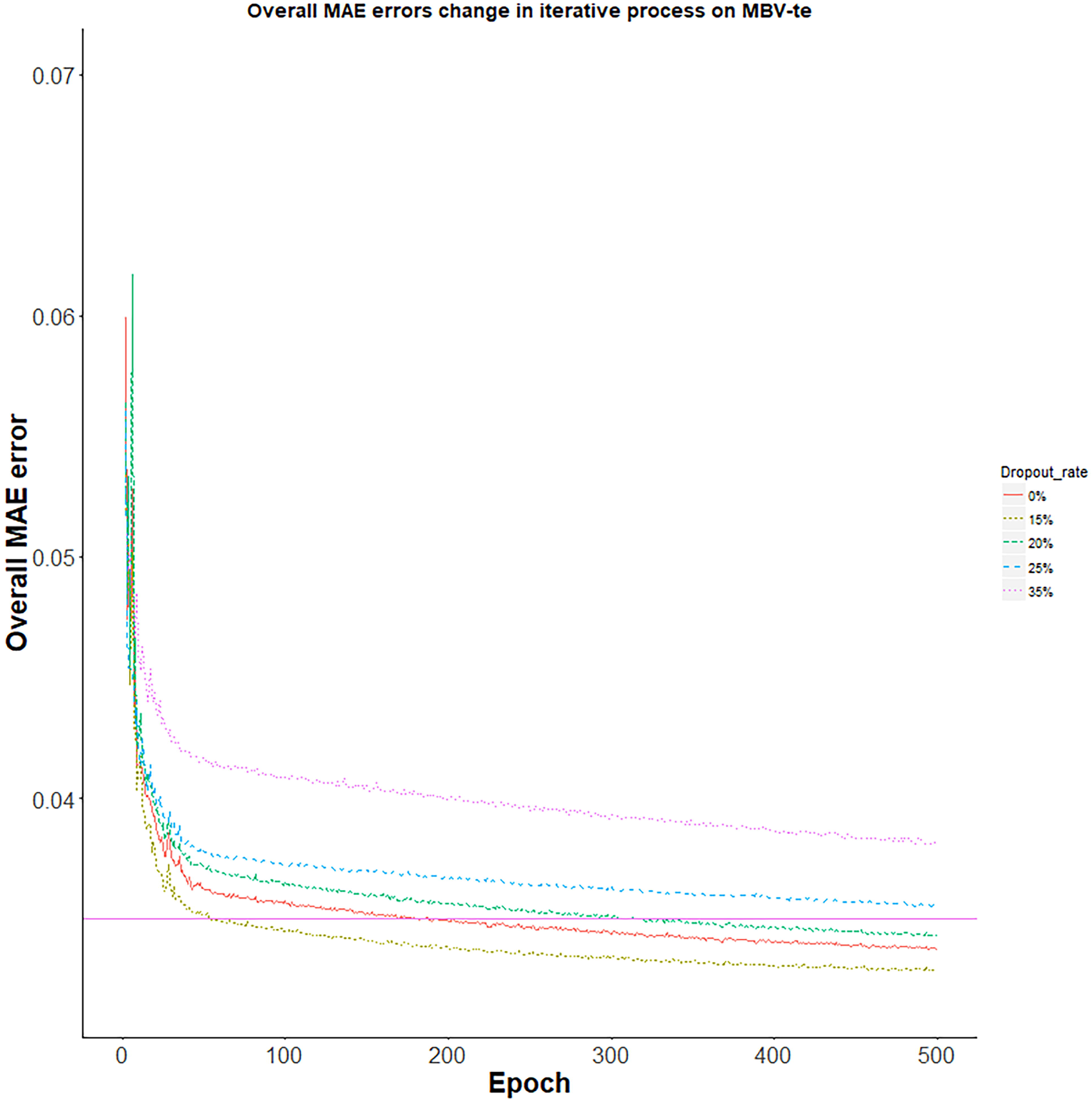
The architecture of D-GPM. It is comprised of one input layer, one or multiple hidden layers and one output layer. All the hidden layers have the same number of hidden units.

## 3 Discussion

Comprehending intricate regulation modes of promoter methylation profiling under numerous biological states needs robust inference frameworks and cost-effective profiling tools. Despite the fact that whole genome bisulfite sequencing is thought of as the best protocol, it is too costly because the genome-wide deep sequencing of the bisulfite-treated fragments needs to be implemented.

Promoter methylation of a gene has been confirmed to be associated with DNA accessibility and the binding of transcription factors that regulate gene expression. Considering that there is an underlying relationship between ~1,000 landmark genes and ~21,000 target genes at the genome-wide level of expression and the fact that methylation occurring in the promoters located in the promoter region is negatively associated with expression of its corresponding gene, we can make good use of the methylation levels in the promoter regions of landmark genes to characterize the cellular status of samples under diverse conditions. Here, we also develop three machine learning methods, namely, LR, SVM and RT, and a deep method called D-GPM for inferring the promoter methylation of target genes based on only landmark genes.

The different methods have different advantages and disadvantages. RT provides us with good interpretability and shows a relative lousy accuracy. LR offers a better performance compared to RT, even though it ignores the nonlinearity within biological data. SVM represents complex nonlinear patterns. Unfortunately, it suffers from poor scalability of the large data size. To some extent, D-GPM manages to overcome the above drawbacks. These prediction models are not perfect and behave better separately for different target genes. It is instructive to interpret these prediction models and explain the inherent relationship between the promoter methylation of the target genes and landmark genes according to our results.

Although D-GPM is the best model for predicting most of the target genes, three machine learning methods all have specific advantages when predicting some specific target genes. Next, we will integrate multiple prediction models as an ensemble tool to ensure it is suitable for predicting as many target genes as possible. In addition, we need to conduct a verification experiment to judge whether the conclusion drawn from the relationship among the promoter methylation of the target and landmark genes (such as *TDP1* and *CIAPIN1*) is sound and persuasive. Furthermore, after obtaining the ensemble prediction model, we will make the most of it to impute and revise methylation sites that are missing or have a low reliability in realistic methylation profiling data.

## 4 Conclusions

In summary, D-GPM acquires the least overall MAE and the highest PCC on the MBV-te among LR, RT, and SVM. For a genewise comparative analysis between D-GPM and the above methods, D-GPM outperforms LR, RT, and SVM in an overwhelming majority of the targets genes, concerning the MAE and PCC. In addition, according to the model distribution of the least MAE and the highest PCC for the target genes, D-GPM predominates among the other models for predicting a majority of the target genes, laying a solid foundation for explaining the inherent relationship between the promoter methylation of target genes and landmark genes via interpreting results from these prediction models.

## 5 Methods

In this section, we first specify the gene methylation datasets in this study and formulate the gene promoter methylation inference problem. We then propose the D-GPM for this problem and describe the relevant details. Finally, we introduce several machine learning methods, which serve as benchmarks.

### 5.1 Datasets

The MBV datasets are acquired from TCGA [23]. Considering that the Illumina

Human Methylation 450k possess more probes and a higher coverage rate, we exclude datasets from the Illumina Human Methylation 27k, and finally, 9756 records remain for the later analysis [24]. After filtering out the records, we calculate the average beta value of all the probes located in the promoter regions of a certain gene as its promoter methylation level. For more information on the data preprocessing, please refer to the data preprocessing section in the Supplementary Material.

We randomly partitioned the methylation data into 80% for training, 10% for validation, and 10% for testing, which corresponded to 7,549 samples, 943 samples, and 943 samples, respectively. We denoted them as MBV-tr, MBV-va, and MBV-te in order.

### 5.2 Multitask regression model for gene expression inference

In the model, there are *J* landmark genes, *K* target genes and *N* training samples. We denoted the training data as 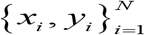, where 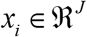 is the promoter methylation profiles of the landmark genes and 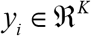 represents the methylation profiles of the target genes in the *i* th sample. Our task is to find a mapping 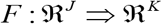 that fits 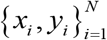 well, which can be viewed as a multitask regression problem.

Let us assume a sample of 9756 individuals, each represented by a 902-dimensional input vector and a 21645-dimensional output vector. Let **X** denote the *N* × *J* input matrix, whose column corresponds to the observations for the *j*-th input 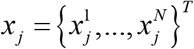. In genetic association mapping, each element 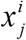 of the input matrix takes values from {0,1,2,3} according to the number of minor alleles at the *j*-th locus of the *i*-th individual. Let **Y** denote the *N* × *K* output matrix, whose column is a vector of observations for the *k*-th output 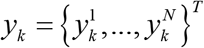. For each of the K output variables, we assume a linear regression model:

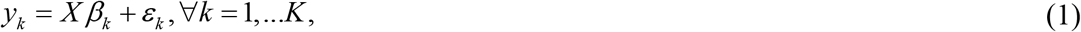

where *β_k_* is a vector of the *J* regression coefficients 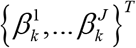 for the *k*-th output, and *ε_k_* is a vector of *N* independent error terms having a mean of 0 and a constant variance. We center the *y_k_’S*,and *x_j_’s*, such that 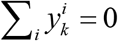 and 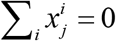, and consider the model without an intercept.

The regression coefficients matrix *β* was used to take advantage of the relatedness across all the input variables.

### 5.3 Assessment criteria

We adopted MAE and PCC as the criteria to evaluate models’ performance at each target gene *t* of the different samples. We formulated the overall error as the average MAE over all the target genes. The PCC was used to describe the relationship between the real promoter methylation and the predicted promoter methylation. Here, the definitions of MAE and PCC for evaluating the predictive performance at each target gene *t* are as follows:

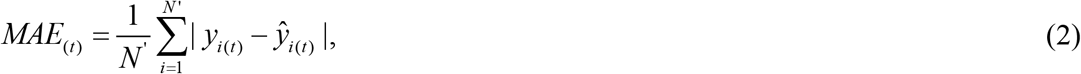

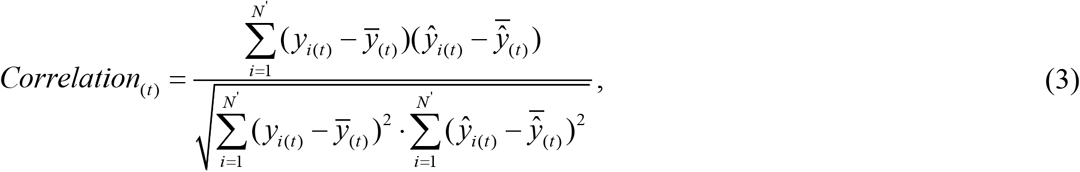

where *N’* is the number of testing samples, 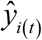 is the predicted expression value for the target gene *t* in sample *i*, and 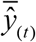 is the mean predicted expression value for the target gene *t* in *N*’ testing samples.

### 5.4 D-GPM

D-GPM is a fully connected multilayer perceptron with one output layer. All the hidden layers consist of H hidden units. In this work, we had a set of Hs, ranging from 1000 to 9000, with a step size of 1000. A hidden unit *j* in layer *l* takes the sum of the weighted outputs plus the bias from the previous layer *l*-1 as the input and produces a single output 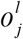.

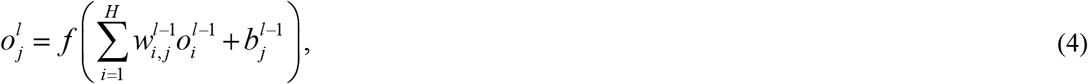

where *f* is a nonlinear activation function, *H* is the number of hidden units, and 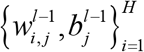 are the weights and the bias of unit *j* to be found.

The loss function is the sum of the mean squared error at each output unit, which is:

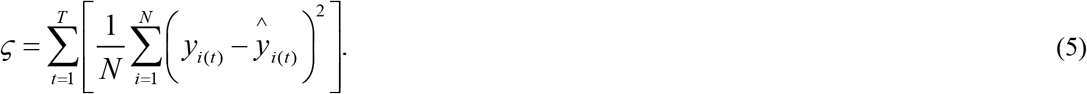

D-GPM contains 902 units in the input layer, corresponding to the 902 landmark genes, and we also configured D-GPM with 21,645 units in the output layer analogous to the 21,645 target genes. Fig. 9 shows the various architectures of D-GPM.

Here, we briefly describe the training techniques and their significance in training steps:

1. Dropout is a scheme used to perform model averaging and regularization for deep neural networks [25]. Here, we utilize dropout to all the hidden layers of D-GPM, and the dropout rate *p*, which steers the regularization intensity, is set at [from 0% to 50%, with step size 5%], separately, to find the optimum architecture of D-GPM.
2. A normalized initialization can stabilize the variances of activation during epochs. To initialize the parameters of deep neural networks, here, we set the initialized weights to within the range of [−1 × 10^−4^, 1 × 10^−4^], according to the activation function.
3. The momentum method is also adopted in our work to speed the gradient optimization and improve the convergence rate of the deep neural networks [26].
4. The learning rate is initialized to 5×10^−4^, 2×10^−4^, 1×10^−4^ or 8×10^−5^, depending on different architecture of D-GPM, and is tuned according to the training error on a subset of MBV-tr.
5. Model selection is implemented based on the MBV-va. The models are assessed on MBV-va after each epoch, and the model with the minimum loss function is saved. The maximum epoch for training is set as 500 epochs.

Here, we implement D-GPM with the Theano and Pylearn2 libraries [27,28].

### 5.5 Benchmark methods

To evaluate the performance of the deep learning methods, we adopt LR, RT, and SVM as benchmarks.

Here, we utilize the RT model with rpart [29]. The Gaussian RBF kernel function has a high superiority for a large sample and for high dimension data, and it reduces the computational complexity in our methylation profiling data efficiently [30];, we adopt the kernlab package to implement SVM for predicting promoter methylation [31].

When training the above three machine models, we harness the 5-fold cross-validation method. Our models are modeled using 80% of the randomly sliced data, and the remaining 20% of the data re used for evaluation. After the process of training and evaluating the model is repeated five times independently, we calculate the average performance during these five processes as a final performance index.

**Abbreviations**
MAE: mean absolute deviation
PCC: Pearson correlation coefficient
LR: linear regression
RT: regression tree
SVM: support vector machine
MBV: methylation beta value
SD: standard deviation

## Supporting information

## Declarations

### Ethics approval and consent to participate

Not applicable.

### Consent for publication

Not applicable.

### Availability of data and materials

The datasets used and/or analyzed during the current study are available from the corresponding author upon reasonable request.

### Funding

This research was supported by the Shenzhen Key Laboratory of Forensics (NO.

ZDSYS201507301424148) of the Shenzhen Municipal Government of China.

### Authors’ contributions

S.C.L. supervised the research and revised the manuscript; X.X.P. designed the study, did the modeling, performed the experiments, carried out the data analyses, graphing and wrote the manuscript; B.L. collected and prepossessed data; X.Z.W. participated in SVM, graphing, manuscript writing and revision; X.X.P., B.L., X.Z.W., Y.L.L., X.Q.Z., and S.B.L. participated in the literature review; all the authors contributed to the manuscript.

## Acknowledgements

The authors acknowledge BGI-Shenzhen for the computing resources and TCGA project organizers for the public datasets.

