## Supplementary material for "D-GPM: a deep learning method for gene promoter methylation inference": Data pre-processing.docx

**1 Data pre-processing**

First, we filtered MBV datasets from TCGA and then cleaned data, including normalization. The landmark and target genes were screened by the LINCS program [1]. Reference TSS is based on hg38. Afterwards, we extracted sample information, including chromosome position, TSS initial position of sense chain, TSS initial position of antisense chain, RefSeqID and gene name from the reference TSS file.

After accessing location information on promoter region of all genes, namely 1500bp from upstream of the TSS site (5'end) to downstream (3'end) 500bp range, we determined relationship among probes and gene promoter regions [2]. After that, the whole methylation value for each promoter region of each sample could be calculated.

Finally, we shuffled our sample data and divide them into training dataset, validation dataset and testing dataset according to the ratio of 8:1:1 for training D-GPM.

1. Edgar, R., Domrachev, M. & Lash, A. E. (2002) Gene Expression Omnibus: NCBI gene expression and hybridization array data repository, *Nucleic acids research.* **30**, 207-10.

2. Schmidhuber, J., Meier, U. & Ciresan, D. (2012) Multi-column deep neural networks for image classification*.* **157**, 3642-3649.
